# Lipidomics analysis of juveniles’ blue mussels (*Mytilus edulis* L. 1758), a key economic and ecological species

**DOI:** 10.1101/768937

**Authors:** Vincenzo Alessandro Laudicella, Christine Beveridge, Stefano Carboni, Sofia Cota Franco, Mary K. Doherty, Nina Long, Elaine Mitchell, Michelle S. Stanley, Philip D. Whitfield, Adam D. Hughes

**Affiliations:** Scottish Association for Marine Sciences, Dunstaffnage Marine Laboratory, PA37 1QA, Oban, UK; Division of Biomedical Sciences, University of the Highlands and Islands, Centre for Health Sciences, Old Perth Road, IV2 3JH, Inverness, UK; Institute of Aquaculture, Faculty of Natural Sciences, University of Stirling, Pathfoot Building, FK9 4LA, Stirling, UK

## Abstract

Blue mussels (*Mytilus edulis* L.) are important components of coastal ecosystems functioning through benthopelagic coupling and ecosystem engineering. At the same time, mussel production is central in the economy of coastal areas. Therefore, understanding their nutritional, physiological and metabolic processes at key life stages is important for their management, both within food production systems and in wild populations.

Lipids are crucial molecules for bivalve growth, but their diversity and roles have been considered from fatty acid (FA) perspective. In this paper, we applied lipidomics to bivalve nutrition. Lipidomics provides a holistic perspective on lipid patterns; by examining the lipidome, important physiological information can be acquired. Here, we use controlled laboratory experiments to elucidate the responses to changes in the diet of newly settled mussels juveniles, one of the most critical life stages. The diets considered in this study are single strains diet of *Cylindrotheca fusiformis* CCAP 1017/2 – CYL, *Isochrysis galbana* CCAP 927/1– ISO, *Monodopsis subterranean* CCAP 848/1 – MONO, *Nannochloropsis oceanica* CCAP 849/10– NANNO and a commercial algae paste –SP.

The diets had a significant effect on spat GR and WI, and according to their efficacy resulted ranked as follows: ISO>NANNO/CYL>SP>MONO. Spat FA composition and neutral lipid content (principally triacylglycerols - TG), were influenced by the diets. Furthermore, untargeted lipidomics also showed shifts in several phospholipid species, with changes related to the essential PUFA available from the diet. TG content, neutral lipids and several TG and FA species were correlated (Spearman R^2^>0.8 FDR p<0.05) with spat WI, suggesting their possible application as markers of mussel juvenile condition. The availability of dietary essential PUFA deeply modified the spat lipidome both for neutral and for polar lipids. This change in the lipidome could have major impacts on their ecology and their production for food.

## 1 Introduction

Blue mussels (*Mytilus* sp.) are ecologically and economically important species, providing a range of crucial ecosystem services along with playing an important role in the economy of many rural and coastal regions [1]. The nutritional value of bivalves is also well documented, providing a source of protein, amino acids, vitamins, trace metals and poly-unsaturated fatty acids – PUFA [2, 3]. Globally, bivalve production accounted for over 15.5% of total aquaculture production in 2016 [4]. Being passive feeders, bivalves reduce the nutrient load in the water [5] whilst not requiring the use of feeds for growth (as observed in intensive culturing of finfish and shrimps [6]), such characteristic makes bivalve culture an environmentally sustainable solution to future food security scenarios [4, 7–9]. Beside their role in nutrients’ bioassimilation, mussels act as a filter for viruses, bacteria, detritus and phytoplankton [10], and as ecosystem engineers, creating shelter and substrate for other benthic organisms and increasing spatial complexity and biodiversity [11]. Furthermore, mussels are classic model organisms in ecotoxicological studies due to their nature of being sessile nature, their ubiquitous presence in the marine environemtn and their filter-feeding behaviour mechanism [12, 13].

European mussel production relies uniquely on natural recruitment events, defined “spatfalls”. Yet, due to influences of environmental drivers [14–16], spatfalls are subjected to severe yearly fluctuations. Such Irregular recruitment, alongside with spreading of diseases, water quality classification and site licensing, is considered between the factors that are preventing the expansion of European mussel production [17, 18]. The establishment of commercial mussel hatcheries is a way to overcome some of these issues. Hatcheries can provide a continuous supply of juveniles to growers, resolving spatfall issues [19]. Nursery operations are a key stage for hatchery production of mussels, as the survival of mussel seedlings on-ropes depends on the size of spat at seeding stage [20–23]. Rearing newly settled mussel postlarvae in nurseries has significant effects on their survival when seeded on grow-out ropes, as it permits to reach a minimal size that ensures higher resistance towards the external environment [19]. However, feeding large amounts of spat in nurseries is expensive, particularly in term of diet supply, as algae production alone accounts around 40% of hatchery costs for rearing bivalve juveniles [24]. Furthermore, juvenile stages are characterised by high production losses, which happen when the hatchery products have their greatest value. Therefore, elucidating the physiological and nutritional requirmentas of mussel juveniles gain a great importance for the development of industrial hatchery production of mussels and of mussel aquaculture production.

Early studies recognised lipids to be the main energy stores for bivalves up to 6 months post-settlement [25, 26]; Since then, evaluating the lipid composition of diets and juveniles gained a central importance on bivalve nutrition studies with examples available for clams [27–33], scallops [34–36], oysters [37–42] and mussels [19, 43–46]. Other than protein and carbohydrate composition, the nutritional properties of shellfish diets strongly depend on the essential PUFA (arachidonic acid – 20:4n-6, AA; eicosapentaenoic acid – 20:5n-3, EPA; docosahexaenoic acid – 22:6n-3, DHA) content [47]. In spite of their ability to *de novo* synthesize lipids, bivalves have limited ability to elongate and desaturate C18 fatty acids (FA) to essential PUFA [27, 29, 48, 49]. FA desaturation requires energy; which could, in turn, penalise growth [30]. To date, a large part of the knowledge on lipids composition and roles in bivalve juvenile nutrition is based on FA analysis. However, this technique only captures a proportion of the lipidome, and lipids have a key role in organism physiology, ranging from biological membranes to cell signalling, immune responses and energy reserves [50]. Current advances in technologies and analytical platforms allow for a deeper and global analysis of lipid molecular species, known as lipidomics. Lipidomics, a branch of metabolomics, encompasses the totality of biological lipids (the lipidome) of a living organism [51]. Lipidomics is mainly based on liquid chromatography coupled with mass spectrometry (LC-MS) platforms [52]. By working with liquid chromatography, information are obtained at lipid molecular species level, rather than on lipid sub-fractions (e.g. fatty acid) or lipid classes. By lipidomics, global changes on lipidome can be visualised, obtaining essential physiological information of the examined organisms [53].

The aim of this paper is to expand existing knowledge on lipid metabolism during nursery operation on mussel juveniles. As such, we applied a comprehensive lipid analysis strategy, which included FA profiling, lipid class analysis and untargeted lipidomics, to the evaluation of the effects of four single strain microalgae species and one commercial algae paste on newly settled *M. edulis* L. 1758. To our knowledge, this is the first time that such holistic lipid analysis approach is applied to bivalve juveniles.

## 2 Materials and Methods

### 2.1 Spat collection and experimental design

Newly settled *M. edulis* (shell length <10 mm) were obtained from Inverlussa Marine Services (www.inverlussa.com; the Isle of Mull, West Coast of Scotland) in July 2017. Spat were collected via gentle scraping of spat collectors, bagged in plastic bags filled with seawater and transferred in ice to the aquarium facilities of SAMS. Upon arrival, the juveniles were graded onto a 4 mm mesh and then kept for 48 hrs on sand-filtered (Grade 1, EcoPure, Waterco) seawater, at room temperature with no food, to allow acclimation and depuration of the animals. After the period of acclimation, spat were sorted and divided into the experiment groups (shell length <5 mm). Groups of 10 selected spat were photographed on milli-graph paper (to obtain shell length – SL) and weighted to 0.0001 g (total live weight – LW). Each group of 10 individuals was successively placed into a 1.5 mm nylon mesh. Three groups of 10 spat constituted the experimental unit (N=30). Three further lots of spat were deployed in an open water environment (OUT) and used as a reference for the laboratory feeding trial.

The feeding trial lasted for 4 weeks, during which the spat were placed in 8l conical tanks, kept at 18 °C in a static system with 20 µm filtered seawater and gentle aeration, under an 18:6 (L:D) photoperiod, with each diet treatment replicated in three independent tanks. The water was fully changed three times per week, coinciding with the feeding of the spat. PH (HI98190, Hanna instruments), salinity (hand refractometer), dissolved oxygen (Fibre optic oxygen transmitter, PreSens) and ammonia (Tetra test NH_3_/NH_4_^+^) were monitored before every water change for the entire feeding trial. Temperature loggers (Pendant, HoboWare) were used to monitor continuously the temperature profile.

Five diet treatments. chosen to have a varied PUFA composition, were evaluated during the trial, one of which included Shellfish Paste (SP – Instant Algae 1800, Reed Mariculture), which is a mixture of *Isochrysis* spp., *Pavlova* spp., *Tetraselmis* spp., *Chaetoceros calcitrans*, *Thalassiosira weissflogii* and *Thalassiosira pseudonana*. The remaining treatments included the administration of microalgae mono-diets of *Cylindrotheca fusiformis* (CYL – CCAP 1017/2), *Isochrysis galbana* (ISO - CCAP 927/1), *Monodopsis subterranean* (MONO – CCAP-848/1) and *Nannochloropsis oceanica* (NANNO – CCAP 849/10). All strains used in this study were provided by the Culture and Collection of Algae and Protozoans (CCAP, www.ccap.ac.uk). Diets were supplied with a weekly ration of 0.4 mg of diet dry weight for each mg of live weight of reared spat [24]. Every week the spat were weighted and their live weight used to calculate the amount of diet to be provided during the following week. The grazing regime was monitored via a turbidimeter (TN-100, Eutech) both before and after diet administration, as an indicator of active grazing (**S1 Fig**). At the end of the trial, both OUT and laboratory kept spat were left for 48 hrs to depurate in filtered seawater and then snap-frozen in liquid nitrogen and stored at −80°C for further analysis. Spat were then freeze-dried (ẞ18 LO plus, Christ) and ground to a fine powder in liquid nitrogen. Ash content was calculated following the combustion of powdered spat for 12 hrs at 450°C.

### 2.2 Microalgae production and their fatty acid composition

Microalgae were grown in sterile 2l Duran’s fitted with aeration lids, tubing and filters kept at 21°C under a 16:8 L:D photoperiod. Media used to maintain each strain are reported in ***Table 1***. To obtain the cell dry weight for each strain (needed to calculate the required weekly food ratio for the spat), 12 aliquots of 1 ml were collected from each stock culture; 6 of them counted using a coulter-counter (Multisizer 3, Coulter Counter) and the remaining 6 freeze-dried to obtain the dry weight which was then reported to the number of cells contained. Weekly, microalgae were harvested via centrifugation at 13G (9000 rpm) for 20 mins 4°C using sterile 250 ml centrifuge tubes (VWR) and concentrated in a 50 ml sterile tubes (VWR) which were kept at 4°C and used for feeding the spat. At every harvesting day, an aliquot (50 ml) from each strain was collected, spun down (14000 rpm 4°C 10 min) and placed in 1.5 ml tube (Eppendorf). The tubes were snap-frozen, freeze-dried and kept at −80°C for lipid analysis.

**Table 1:**
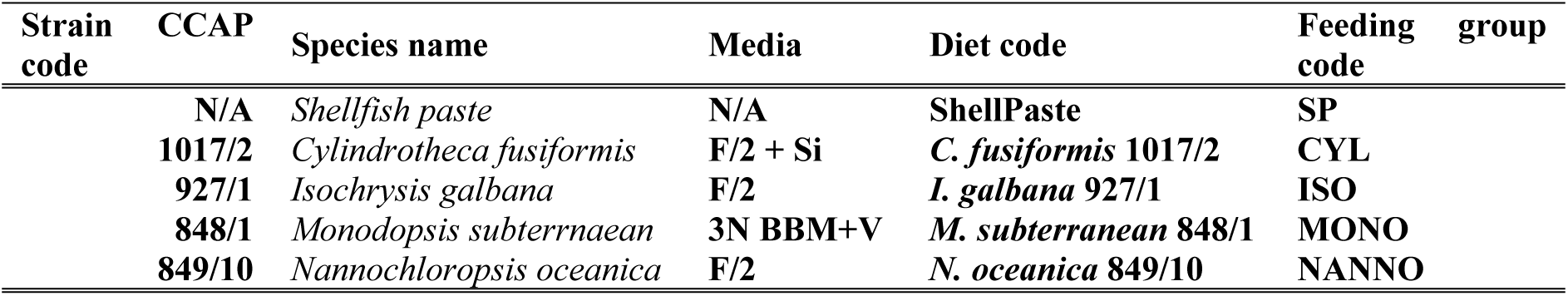
Summary of diets employed during the feeding trial. For the live algae treatments, media used in the culture of microalgae strains and relative CCAP codes are reported.

The relative (%FAME) and absolute (μgFA mg_DW_^−1^) composition of the diets employed in the trial is reported in **Table 2**. Total saturated FA (SAFA) was highest in *C. fusiformis* and *I. galbana* (p<0.05), while the highest amount of monounsaturated FA (MUFA) was observed in *N. oceanica. C. fusiformis* resulted in the richest diet for n-6 PUFA (p<0.001) whilst no evident differences between diets were found for total n-3 and total PUFA content (p>0.05). ANOSIM (R 0.92 p<0.001) evidenced the presence of multivariate differences between the various diet. *I. galbana* was characterised by a high content of 22C FA as 22:4n-6, 22:5n-3 and DHA (p<0.001), while lacked EPA and AA. Further relevant FA in *I. galbana* resulted 14:0, 18C FA as 18:1n-9, 18:2n-6, 18:3n-3 and 20:1n-11. On the contrary *C. fusiformis* and *N. oceanica*, respectively resulted rich in AA (p<0.001) and EPA (more abundant in *N. oceanica*, p<0.01) and the MUFA 16:1n-7, which accounted for the 20% of total FA in both strains. *M. subterranean* was characterised by the richness of C18 FA as 18:2n-6, 18:3n-6 and 18:3n-3 (p<0.001). *N. oceanica* and *C. fusiformis* were poor sources of 18:4n-3, which accounted for the 5% of total FA in all the other diets (p<0.001). ShellPaste, as a mixture of different microalgae strains, resulted in a balanced composition of main essential PUFA as EPA (16.1%) and DHA (6.17%), while lacked in AA (0.46%).

**Table 2:**
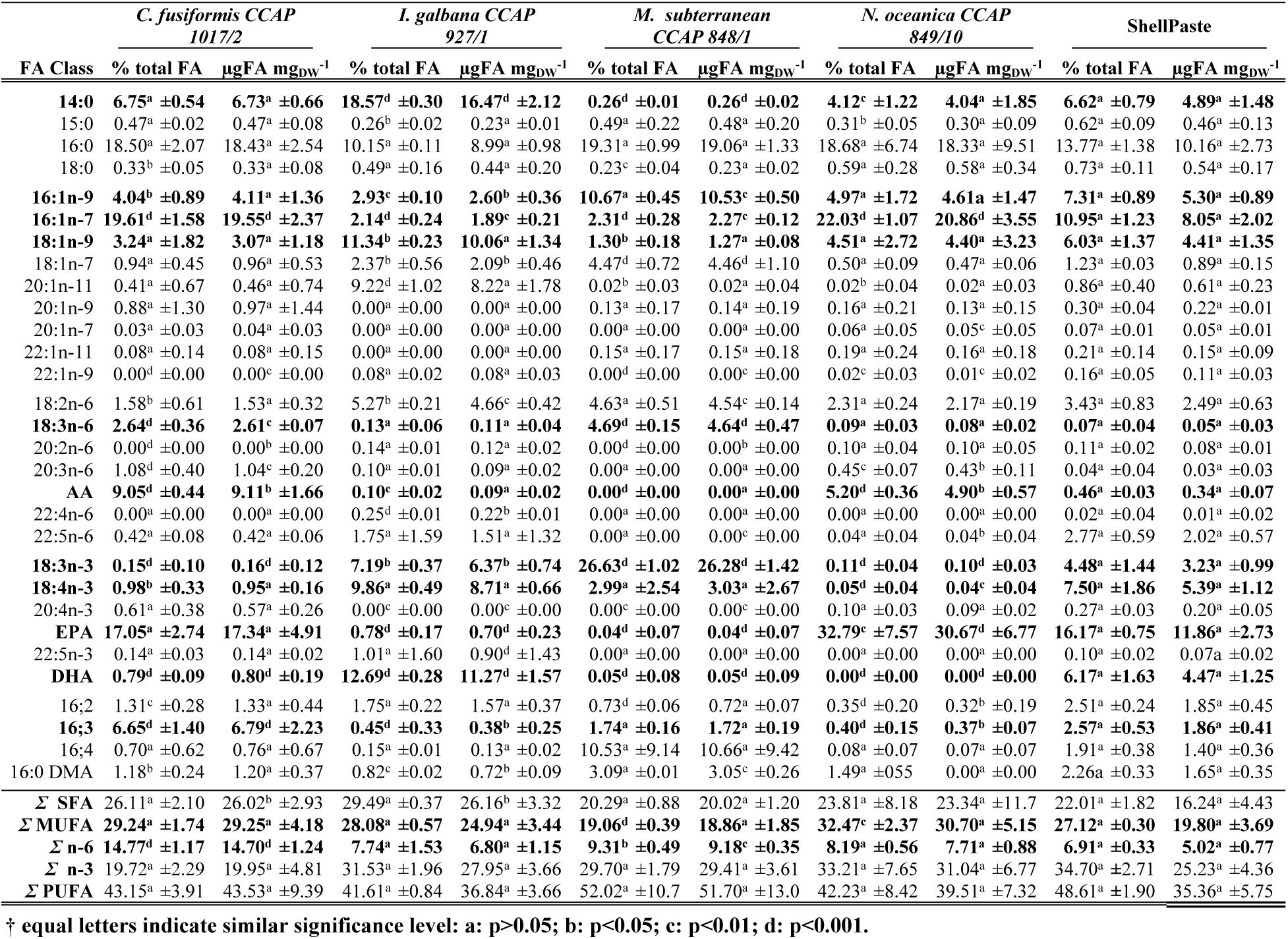
Fatty acids composition of the five diets employed in this study, reported as percentage of each FAME for the total fatty acid content for each diet (% total FA) and as absolute fame content µg of FA per mg of algae dry weight (µgFA mg_DW_^−1^). AA: arachidonic acid – 20:4n-6, EPA: eicosapentaenoic acid – 20:5n-3, DHA: docosahexaenoic acid – 22:6n-3; DMA: dimethylacetals. Data are reported as average of three replicates ± SD. Statistical differences are reported in comparison to ShellPaste. FA evidenced by SIMPER and differing significantly between diets are in **bold**. Letters correspond to statistical significance: a p>0.05, b p<0.05, c p<0.01, d p<0.001 (**†**).

### 2.3 Biometrical analyses

Spat growth performances (GR) was measured as shell length increase (SI) and live weight increase (WI). At the beginning and at the end of the trial, spat were photographed on milli-graph paper, and their SL was obtained by processing the images via ImageJ software (www.imagej.nih.gov). GR was calculated as Δ between shell size at T28 and T0. For WI calculation, spat were blotted on tissue paper and weighted at 0.0001 g scale (Sartorius). The Δ between the live-weight at the beginning of the trial and the end resulted in WI.

### 2.4 Biochemical analyses

#### 2.4.1 Lipid extraction

The homogenization procedures changed according to the matrix. Microalgae and diet samples were resuspended in 200 µl of milliQ water and disrupted via probe sonication for 1 minute, whilst aliquots (≈ 10 mg) of powdered spat were homogenated in 200 µl of milliQ water by pestling in ice for 1 minute. For all samples, lipid extraction was done according to Folch, Lees (54). The dried lipid extracts were weighted to the 0.00001 g (Sartorius) and resuspended in 0.5 ml of chloroform constituting the total lipid extract (TLE). The TLE was divided into 2 sub-aliquots. One aliquot (400µl) was dried down in nitrogen and stored at −80 °C for lipid class and lipidomics analysis. To the second aliquot (100 µl) was added an internal standard (FA 17:0, Sigma + 0.001% of BHT, Cayman Chemical Company at the 10% of the total lipid mass) and processed for fatty acid methyl esters (FAME) analysis.

#### 2.4.2 Fatty acids analysis

FAME from TLE of diets and spat were prepared by acid-catalysed transesterification according to AOCS (55). The FAME layer was evaporated under a gentle nitrogen stream (NVap, Organomation) and the FAME were resuspended in 500 µl of iso-hexane, purified on silica SPE cartridges (Clean-up Cusil 156, UCT) preconditioned with 5 ml of iso-hexane, and eluted twice with 5 ml of a 95:5 iso-hexane:diethyl ether solution. Purified FAME were dried in N-vap and resuspended to 1 mg ml^−1^ according to the original lipid mass in iso-hexane (HPLC grade, Fisher).

FAME were separated by gas chromatography using a ThermoFisher GC 8000 (ThermoScientific) equipped with a fused silica capillary column (ZBWax, 60m x 0.32 x 0.25 mm i.d.; Phenomenex) with hydrogen as a carrier gas and using on-column injection. The temperature gradient was from 50 to 150°C at 40°C min^−1^ and then to 195°C at 1.5°C min^−1^ and finally to 220°C at 2°C min^−1^. Individual methyl esters were identified by comparison to known standards (Marine Oil, Restek) and by reference to published data [56]. Data were collected and processed using the Chromcard for Windows (version 2.00) computer package (Thermoquest Italia S.p.A.). Results are reported as a relative percentage of FAME composition (%FAME) and as absolute quantification via the internal standard method (µglipid mg_DW_^−1^ for microalgae and µgFA mg ^−1^ in the case of spat).

#### 2.4.3 Lipid class analysis

TLE from spat and diets were separated in their main lipid class via normal phase high-pressure liquid chromatography coupled with electron light scattering detector (NP-HPLC-ELSD). The separation was accomplished with an Infinity 1260 platform (Agilent Technologies) according to Graeve and Janssen (57) with minor modifications. The protocol was modified to enhance the separation of certain lipid classes relevant in marine invertebrates, as phosphonoethyl ceramides (CPE). TLE was separated on a monolithic silica column (Chromolith Si 100×4.6, Merck) equipped with the relative guard columns (Chromolith Si guard cartridges, Merck). The column was kept at 40 °C and the solvent flow kept at 1.4 ml min^−1^. The quaternary elution gradient is reported in ***Table 3***. Acetone, Isooctane (tri-methyl pentane), Ethyl Acetate and Water were all HPLC grade and obtained from FisherBrand, HPLC grade isopropanol (IPA) was obtained from Chromanorm. Glacial Acetic Acid (GAA) and Triethylamine (TEA), both HPLC grade, were purchased from VWR.

**Table 3.**
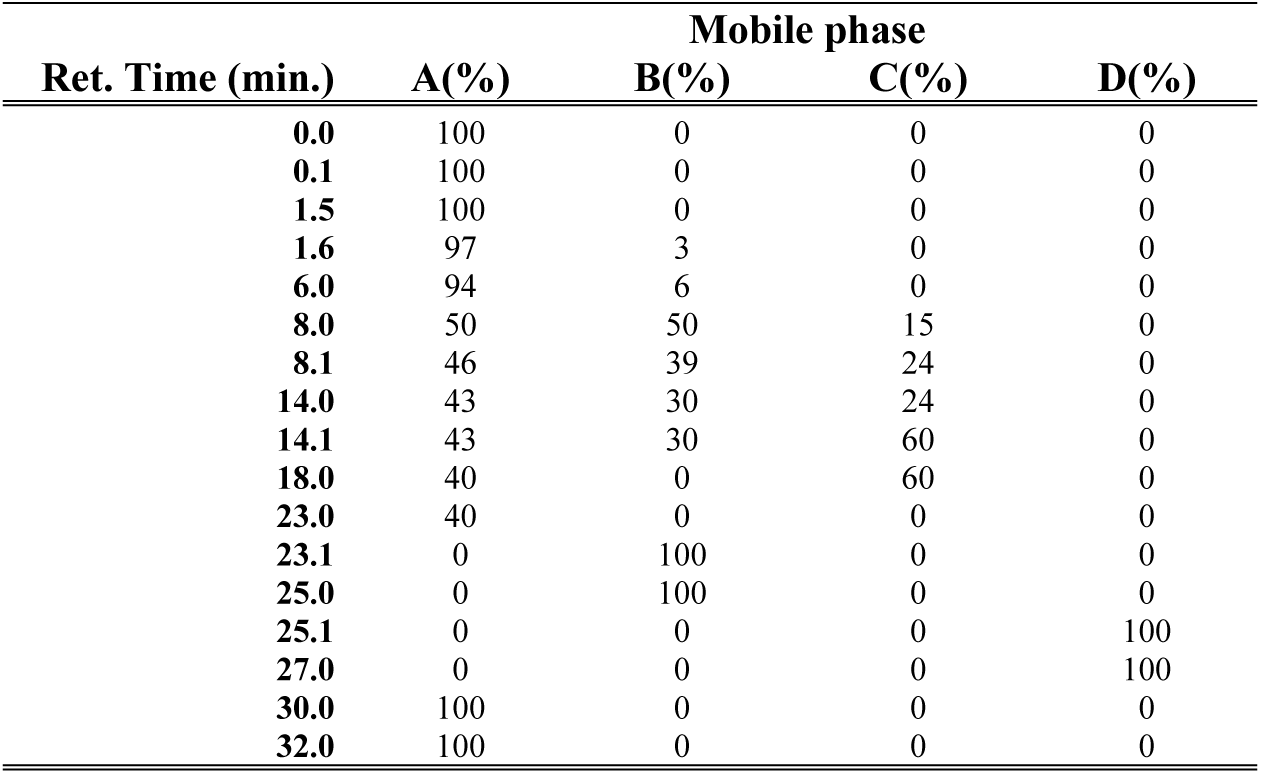
Quaternary gradient used during NP-HPLC separation of spat TLE. Mob A: Isooctane:Ethyl Acetate (99.8:0.2); Mob B: Acetone: Ethyl Acetate (2:1) + 25 mM GAA; Mob C: IPA: Water (85:15) + 15mM GAA and 7.5 mM TEA; Mob D: Isopropanol.

Identification of principal lipid classes was achieved via an external standard method. Commercially available purified lipid fractions were used as lipid class standards. Squalene (terpenes - TER), Arachydil dodecanoate (wax ester – WE), Cholesterol (free sterols – ST), triglycerides (triolein – TG), diacylglycerols (DG), monoacylglycerols (MG), FA17:0 (free fatty acid – FFA), Ceramides lipid mix from bovine brain (Cer), Phosphatidic acid (PA), phosphatidyl ethanolamine from soybean (PE), Cardiolipin from bovine heart (CL), Phosphatidyl serine (PS) and phosphatidyl choline (PC), phosphatidyl inositols (PI), lyso-phosphatidyl choline from egg yolk (LysoPC) all obtained by Sigma, sphingosylphosphorylethanolamine (CPE) from Matreya. Stock solutions for each lipid were prepared at 2.5 mg ml^−1^ in 2:1 chloroform:methanol (HPLC grade, Fisher). From the stock solution, working solution were diluted in Mob A. Calibration curves were calculated by sequential 10 µl injections of standard mix solutions (0.5-0.25-0.125-0.066-0.033-0.0165-0.008 μg µl^−1^ of each lipid class). Identification of lipid classes was achieved by retention time match between unknown samples and the standard mix. Spat TLE were resuspended in a 4:0.06:0.04 (Mob A:chloroform:methanol) solution at a concentration of 1 mg ml^−1^ of which 10 µl volume was injected. Chromatograms were inspected, integrated and calibration curves calculated via Chemstation software (Agilent Technologies). Results are reported as relative lipid class composition (%TLE) and in absolute values (µglipid mg_ash free DW_^−1^).

#### 2.4.4 Untargeted lipidomics

Untargeted lipidomics of spat was achieved via High-Resolution Mass spectrometry (HRMS). The platform used was a binary HPLC (Accela, ThermoFisher) coupled with an electron spray ionization (ESI) and orbitrap mass analyser (Exactive, ThermoFisher). The separation was done on a C18 Hypersyl Gold 100×2.1 mm 1.9nm particle size (ThermoFisher) kept at 50 °C. The binary solvent system included a constant flow rate of 400 μl min^−1^ with a gradient as described in **Table 4**. Water and Acetonitrile were HPLC grade and obtained from Fisher, IPA was LC-MS grade (Hypergrade LiChrosolv, Merck), while ammonium formate and formic acid were both LC-MS grade and obtained from Sigma Aldrich.

**Table 4:** Binary gradient used during LC-MS analysis of spat TLE. Mob A: Water + 10 mM Ammonium formate + 20 mM Formic acid; Mob B: IPA:ACN (9:1) + 10 mM ammonium formate + 20 mM formic acid.

| Ret. Time (min.) | Mobile phase |  |
| --- | --- | --- |
|  | A(%) | B(%) |
| 0.0 | 65 | 35 |
| 0.5 | 65 | 35 |
| 4.0 | 35 | 65 |
| 19.0 | 0 | 100 |
| 21.0 | 0 | 100 |
| 21.1 | 65 | 35 |
| 27.0 | 65 | 35 |

The mass spectra were acquired in the m/z range 250-2000 both in positive ESI (POS) and in negative ESI (NEG) with a mass resolution power of 100,000 FWHM. The mass error was kept below 5 ppm by routinely calibrations on both polarities with a calibration solution (Pierce™ LTQ ESI calibration solutions, ThermoFisher). TLE from spat were resuspended in 3:1 methanol: chloroform at a concentration of 1 mg ml^−1^ with 3 μl injection volume. Chromatograms and mass spectra were inspected and integrated via Xcalibur software (ThermoFisher), data processing and analysis procedures are reported in the data analysis section. LC-MS profiles of spat extracts are reported in **S2 Fig**. Features were identified according to their precursor ion exact mass (MS’) and reported as lipid class with the total number of carbons and double bonds (e.g. PC(36:5), for phosphatidyl choline with 36 carbons and 5 double bonds on the fatty acyl residues). Isobaric lipids separated in reverse phase chromatography but evidencing same MS’, are reported with different letters (e.g. PC(38:5)_a_ PC(38:5)_b_).

### 2.5 Data analysis

#### 2.5.1 Biometrical, FAME and lipid class analyses

Statistical analysis was compiled via R statistical software (version 3.5.1). Data are reported as mean ± standard deviation. Statistical differences were considered significant for p<0.05. Biometrical data were log-transformed to force homoscedasticity. If normality assumptions were met, a two-way analysis of variance (two-way ANOVA) and a Tukey HSD test were employed to evaluate differences between the different diet groups at each sampling point. If homoscedasticity, following data transformation, was not met a Kruskal-Wallis with a Dunn’s test (R ‘dunn.test’ package) as *posthoc* test was used to evaluate the effects of diet treatments on the spat.

Lipid class and FAME data were square-root transformed and multivariate differences were evaluated via Analysis of Similarities (ANOSIM). Non-Metric Multidimensional Scaling (nMDS) with Euclidean distance matrix was employed to graphically plot each sample group. Similarity percentages (SIMPER) were applied to evaluate the main lipid and fatty acids characteristic for each groups clustering. ANOSIM, nMDS and Simper analysis were obtained from R ‘vegan’ package [58]. Lipid class and fame composition differences between groups were evaluated via a one-way ANOVA and Tukey HSD (false discovery rate – FDR – adjusted p-value [59]) as *posthoc* test, whilst Kruskal-Wallis with a Dunn’s test as *a posteriori* comparison was employed on features that failed normality assumptions (tested via a Cochran test).

#### 2.5.2 Untargeted Lipidomics

Raw LC-MS data were processed via Progenesis QI software (Nonlinear Dynamics, Waters). A technical QC sample was run every 6 hours of instrument operation time to monitor possible technical shift in the machine. Chromatograms were automatically aligned using a QC as a reference point. Peak picking and deconvolution were completed following automatic settings of the software, with intensity threshold of 1xE^5^ and 1xE^4^ respectively for POS and NEG ionization modes. Data were normalised according to the total ion current of each chromatogram. Main lipid adducts for both POS and NEG were experimentally evaluated by using a lipid standard mixture that included the main lipid classes and were added to the software search (**S1 Table**). Lipid identification was achieved by searching the lipid dataset versus LIPID MAPS (www.lipidmaps.org), HMDB (www.hmdb.ca), Metlin (www.metlin.scripps.edu) and an “*in house*” bivalve lipid database built from recent bivalve lipidomics studies [60–69]. Contaminants were manually removed from the peak intensity table (PIT) generated from this process.

The PIT was furtherly filtered and processed with the R based package ‘MetaboAnalystR’ [70]. Filtering process included the removal of features with over 30% of missing values and substitution of remaining missing values with a small value (half of the minimum intensity value). Features with low repeatability or low constant values were filtered out using QC samples and inter-quantile range, data was then scaled via Pareto scaling, to reduce the skewness of data and enhance comparability between different samples. Chemometrics analysis was employed as data reduction and biomarker discovery tools. Principal component analysis (PCA) was used to evaluate data quality, clustering between QC samples and the presence of outliers (**S2 Fig**). Partial least squares discriminant analysis (PLS-DA) was applied to cluster samples and to calculate variables of importance in projection (VIP) scores, which represent the weighted sum of squares of the PLS loadings taking into account the amount of explained Y-variation [71]. PLS-DA model fitting was evaluated via a permutation test and ten-fold leave one out cross-validation (LOOCV, **S3-S6 Fig**). The statistical significance of the features with VIP scores >1 was furtherly screened via a Kruskal-Wallis test with FDR adjusted p-value [59].

#### 2.5.3 Identification of main lipids linked with growth spat performances

The correlation between spat growth performances (in term of WI) and spat lipid composition (considering the significant lipids evidenced from multivariate analysis of FAME, lipid class and lipidomics dataset) was calculated by means of Spearman rank correlation coefficient. Absolute values for lipid class and FAME data and relative intensity for lipidomics data were used to compute the correlation.

## 3 Results

The diets employed in this study were selected due to their distinct FA profile (**Table 2**). *I. galbana* presented DHA as main essential PUFA (12.69±0.24% /11.27±1.57 µgFA mg_DW_^−1^). AA and EPA were the most abundant PUFA in *C. fusiformis* and *N. oceanica*, with AA resulting higher in *C. fusiformis* than *N. oceanica* (respectively 9.05±0.44% / 9.11±1.66 µgFA mg_DW_^−1^ and 5.20±0.56% / 4.90±0.57 µgFA mg_DW_^−1^) and with EPA showing the opposite trend (respectively 17.05±2.74% / 17.34±4.91 µgFA mg_DW_^−1^ and 32.79±7.57% / 30.67±6.77 µgFA mg_DW_^−1^). On the other hand, *M. subterranean* lacked in C20 PUFA and was characterised by large amounts of 18:3n-3 (26.63±1.02% / 26.28±1.42 µgFA mg_DW_^−1^). We included ShellPaste on the experimental design to have a diet, composed by more than one microalgae strains, which supplied EPA and DHA (respectively 16.17±0.75% / 11.86±2.73 µgFA mg_DW_^−1^ and 6.17±1.63% / 4.47±1.25 µgFA mg_DW_^−1^).

At the end of the feeding trial period, we observed that the spat GR significantly varied across the sample groups (**1**). After 4 weeks of deployment at sea, OUT resulted the group with the longest shells (shell length – SL, 4.86 ± 0.68 mm) and highest live weight (LW, 75.08 ± 14.2 mg_LW_ spat^−1^). Although smaller than OUT, also spat fed with *I. galbana* (ISO), *N. oceanica* (NANNO), *C. fusiformis* (CYL) and shellPaste (SP) resulted in bigger shells compared with the beginning of the trial (p< 0.001, **Figure1A**), whereas spat fed with *M. subterranean* (MONO) did not present any significant increase in their shell lenght. LW significantly increased in ISO, CYL and SP treatments (Although SP LW resulted lower than the other groups, **Figure 1C**). Nevertheless, if GR is considered (SI and WI, **Figure 1B-D**) ISO and OUT outperformed the remaining sample groups (p<0.01).

**Figure 1:**
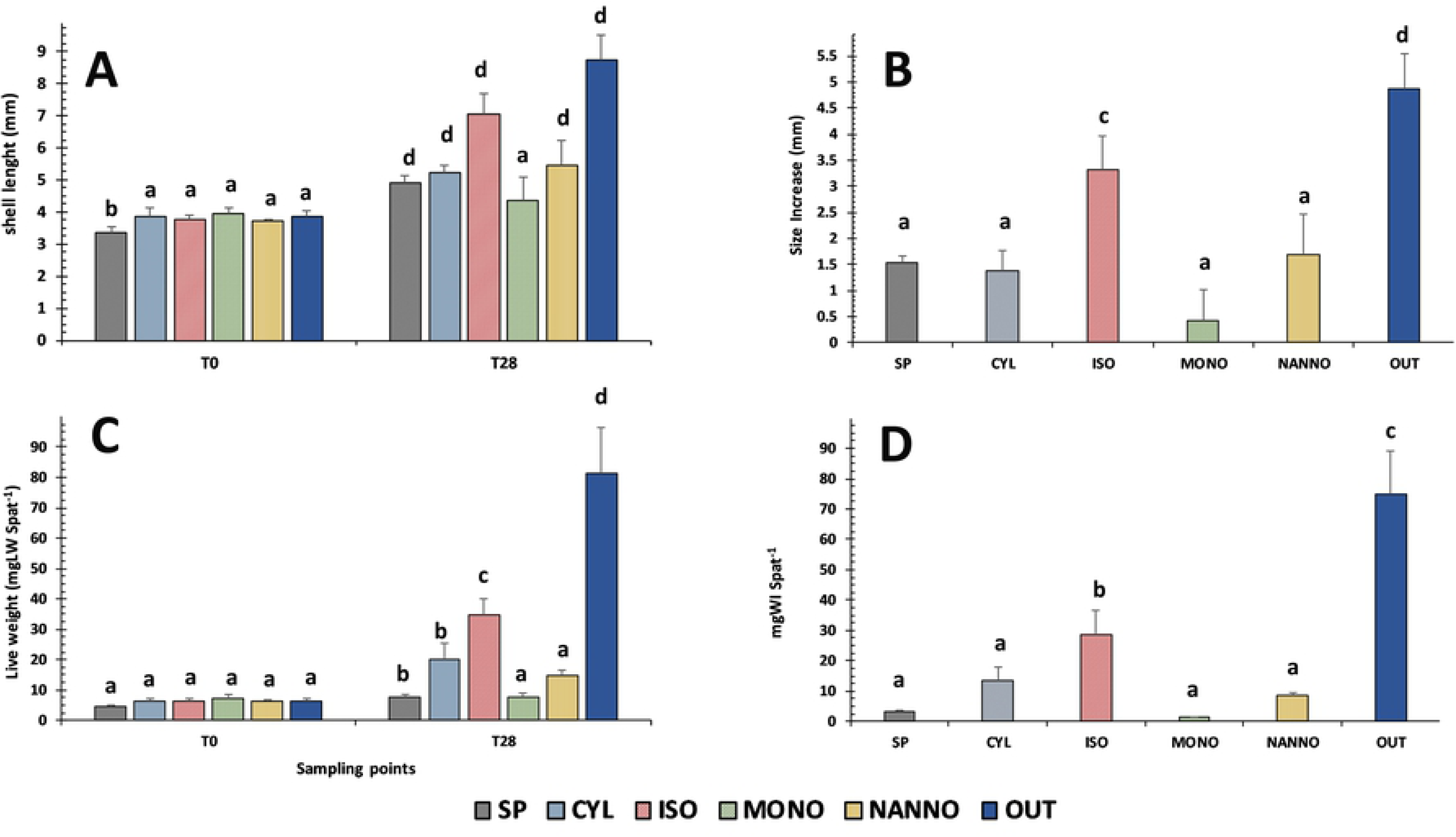
Growth rate (GR) and live weight increase (WI) on different spat groups. A: variation of shell length in mm between in each group between the beginning of the trial (T0) and end of the experiment (T28). B: Size increase of the spat (reported in mm) during the diet trial when subjected to different diets. C: live weight per spat (reported in mg) between the beginning of the trial (T0) and the end of the trial (T28). D: weight increase per spat throughout the trial according to the different sample groups (reported in mg). SP: Shellfish Paste, CYL; *C. fusiformis* 1017/2, ISO: *I. galbana 927/1*, MONO: *M. subterranean 848/1*, NANNO: *N. oceanica 849/10*, OUT: Outdoor. Data are reported as average ± SD. For figure A and C N=30 individual spat; for figures B and D N=3. Letters indicate significance levels: **a: p>0.05; b: p<0.05; c: p<0.01; d p<0.001;**

**Figure 2.**
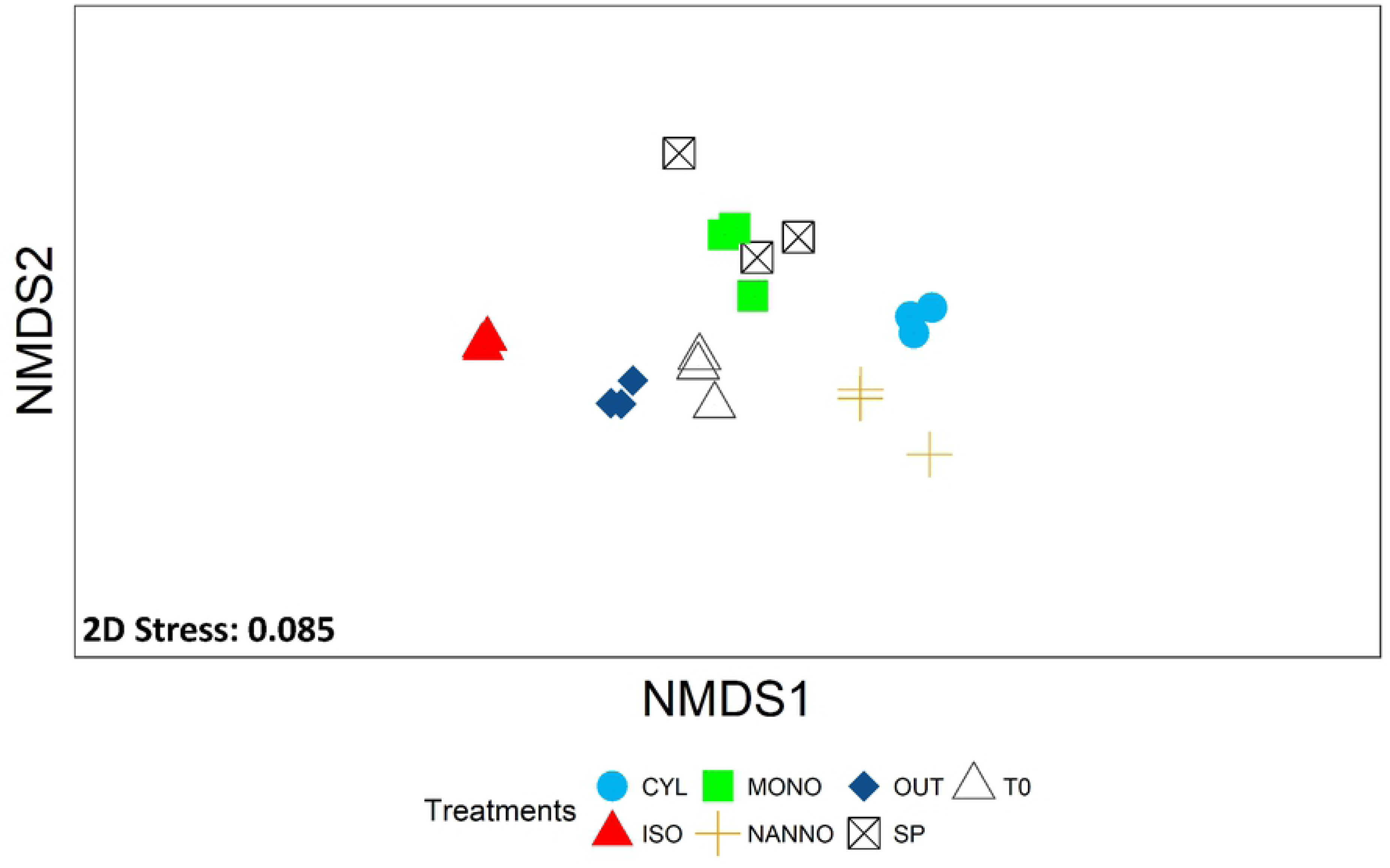
Non-metric multidimensional scaling (nMDS) plot of the relative FAME composition of spat (%FAME) subjected to the feeding trial. T0: Spat sampled before the beginning of the trial, SP: spat fed with Shellfish Paste during the 4 weeks trial; CYL: spat fed with *C. fusiformis* 1017/2 during the 4 weeks diet trial; *ISO:* spat fed with *I. galbana* 927/1 during the 4 weeks diet trial; MONO: spat fed with *M. subterranean* 848/1 during the 4 weeks trial; NANNO: spat fed with *N. oceanica* 849/10 during the 4 weeks trial; OUT: spat deployed outdoor and sampled after 4 weeks. Three replicates (n=3) for each sample group are here reported.

The investigation continued with the evaluation of the effects on the lipidome of the spat, beginning from traditional lipid analysis techniques as FA and lipid classes profiling. The data, expressed as relative composition (%) and absolute value (µg mg ^−1^) of FA and/or lipid classes, are reported in ***Table 5***. ANOSIM analysis suggested the presence of strong differences both for FA and lipid class between sample groups (R 0.935 p <0.001 for FA and R 0.744 p<0.001 for lipid class). During the feeding trial, the absolute amount of saturated fatty acids (SFA) and mono unsaturated fatty acids (MUFA) resulted low in poor performing groups (MONO and SP). Laboratory-reared spat (SP, CYL, ISO, MONO and NANNO) resulted richer their relative total n-6 PUFA compared with wild spat (T0 and OUT, p<0.001), with the highest absolute content observed in CYL and ISO (p<0.05). The relative content of n-3 PUFA significantly decreased in CYL, MONO, NANNO and SP (p<0.001), while was significantly higher in OUT (p<0.01) when compared with T0. The percentage of total PUFA resulted significantly higher in OUT and ISO while it was significantly lower than T0 in MONO and NANNO (p<0.01). The absolute content of these last two parameters decreased significantly in MONO and SP compared with T0 (p<0.05).

**Table 5:**
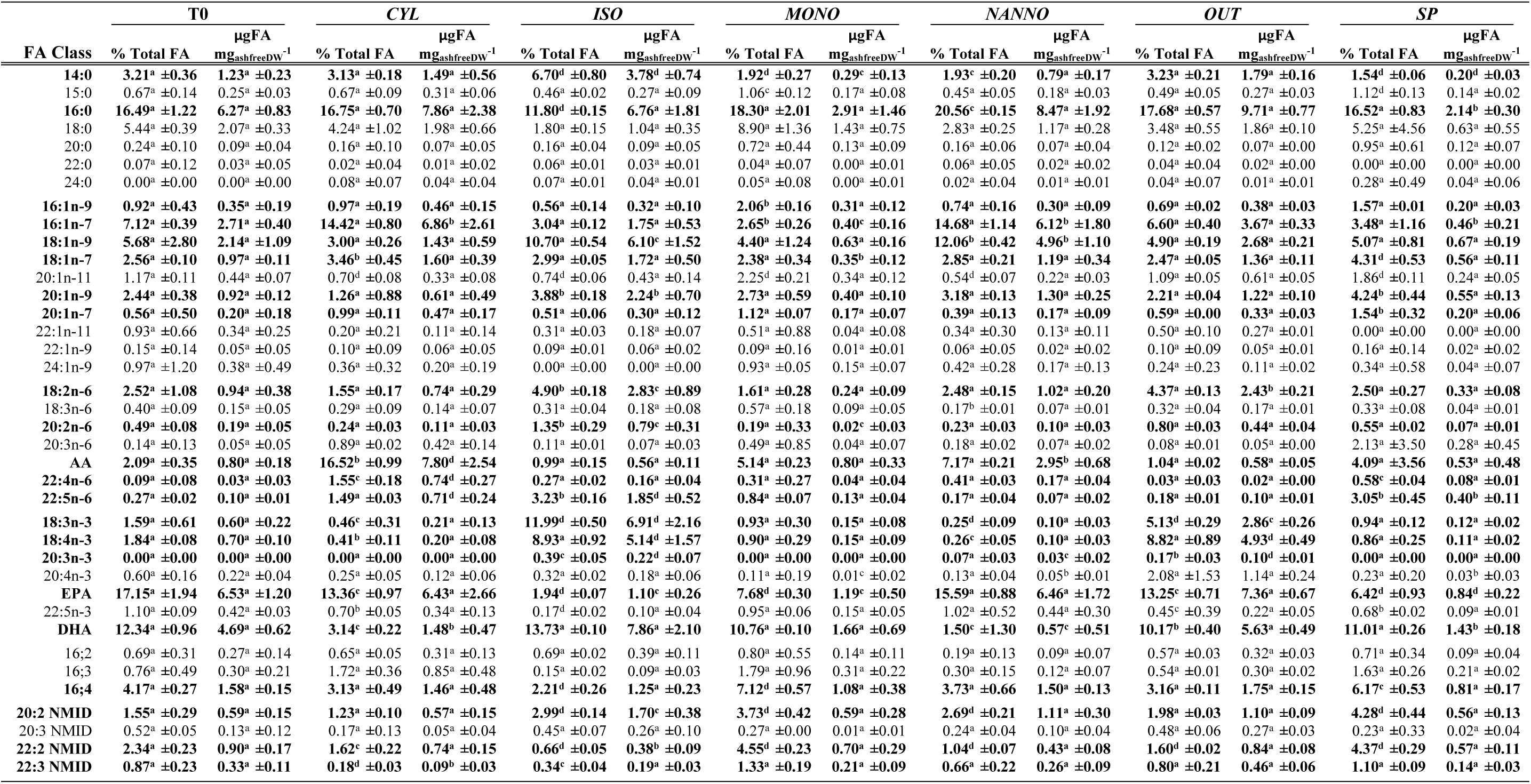
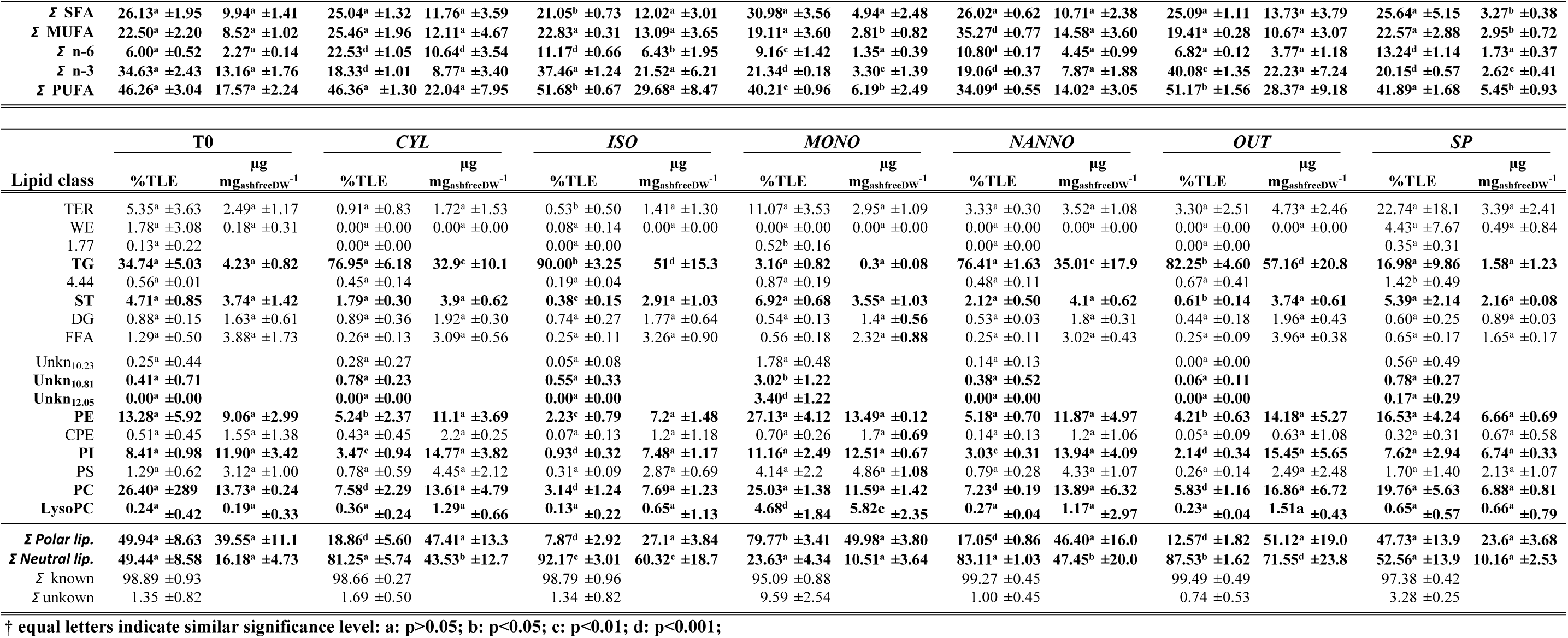
Fatty acids and lipid class composition of the spat subjected to the diet treatments, reported as % of each FAME for the total fatty acid (or of the total lipid extract – TLE) and absolute content (µgFA mg_ashfreeDW_^−1^ or µglipid mg_ashfreeDW_^−1^ spat) for each diet group. FAME acronyms: AA: arachidonic acid – 20:4n-6, EPA: eicosapentaenoic acid – 20:5n-3, DHA: docosahexaenoic acid – 22:6n-3; NMID: non-methylene interrupted dienoic fatty acids. Lipid class acronyms: WE: wax esters, TG: triacylglycerols, ST: free sterols, DG: diacylglycerols, FFA: free fatty acids, PE: phosphatidylethanolamines, CPE: ceramide phosphoethanolamines, PI: phosphatidylinositol, PS: phosphatidylserine, PC: phosphatidylcholine, LysoPC: lysophosphatidylcholine. Unidentified lipid classes are reported with Unkn and their retention time span (e.g. Unkn_12.05_). T0: Spat sampled before the beginning of the trial, SP: spat fed with Shellfish Paste; CYL: spat fed with *C. fusiformis* 1017/2; *ISO:* spat fed with *I. galbana* 927/1; MONO: spat fed with *M. subterranean* 848/1; NANNO: spat fed with *N. oceanica* 849/10; OUT: spat deployed outdoor and sampled after 4 weeks. Data are reported as a average of three replicates ± SD. Statistical significance is reported in comparison with T0. FA and lipid class evidenced by SIMPER and differing significantly between sample groups are in marked in **bold**. Letters correspond to statistical significance: a p>0.05, b p<0.05, c p<0.01, d p<0.001 (**†**).

Further differences between treatment groups were analysed via nMDS and SIMPER. NMDS plot (stress=0.085) is reported in **Error! Reference source not found.**. T0 clustered in the centre of the plot and from there three main clustering group of samples are observed. OUT and ISO closely clustered together, characterised by the content in 18:2n-6 (p<0.05), 18:3n-3 (P<0.01) and 18:4n-3 (p<0.001), 20:3n-3 (p<0.001). ISO was also characterised by high amount of 14:0 (p<0.001), 18:1n-9 (p<0.01), 20:1n-9 (p<0.05), of the PUFAs 20:2n-6 (p<0.05), 22:5n-6 (p<0.05) and 20:3n-3 (p<0.05) and FA20:2 non methylene interrupted dienoic (NMID, p<0.01). EPA was significantly lower in ISO than T0, CYL, NANNO and OUT (p<0.01). CYL and NANNO, clustered closely together on nMDS plot; these groups were characterised by the rich amount of 16:1n-7 (p<0.05), AA (p<0.05) and 22:4n-6 (although only in CYL resulted different from T0, p<0.05) and low DHA content (p<0.01). CYL was also characterised by the presence of 22:4n-6 and 22:5n-6 (p<0.05), whilst higher levels of 18:1n-9 and the relative content of FA 20:2 NMID characterised NANNO (p<0.05). A reduction of FA 22:2 and 22:3 NMID was also observed in CYL and ISO (p<0.05). SP and MONO closely clustered the nMDS plot as characterised by low EPA (p<0.01) and their high relative content of 16:4, 20:2 and 22:2 NMID (p<0.01). DHA content was significantly lower in SP compared with T0 (p<0.05).

The analysis of lipid class showed that the main lipid classes found in spat included terpenes (TER), wax esters (WE), triacylglycerols (TG), free sterols (ST), diacylglycerols (DG), free fatty acids (FFA), phosphatidylethanolamines (PE), phosphatidylinositols (PI), phosphatidylcholines (PC) and lysophosphatidylcholine (LysoPC). The nMDS analysis of spat lipid classes (***Figure 3***) evidenced the presence of three main groups of samples according to the diet provided (stress= 0.028). T0 and SP contained similar relative amounts of neutral and polar lipids and occupied the central portion of the nMDS plot. A large % of neutral lipids in these samples were constituted by terpenes (varied between 5% to over 22% of TLE) and TG (16-30% of TLE). Due to the lower relative abundance of TG, higher relative amounts of polar lipids were observed in these groups (***Table 5***; p<0.05). Neutral lipids dominated the relative composition in ISO, OUT, CYL and NANNO, which clustered closely on the nMDS plot. In all these groups, the main lipid class was triacylglycerols (TG), which accounted for over 75% of TLE in these samples. The predominance of TG in the previous diet groups, consequently affected the relative quantification of the polar lipid classes, resulting in significant differences when compared with T0 (p<0.05). Lastly, MONO clustered isolated from the other samples, characterised by the lowest amount of TG, two unidentified lipids Unkn_11.80_ (p<0.05) and Unkn_12.05_ (p<0.001) and by the relative content in polar lipid (79.77±3.41%, p<0.05).

**Figure 3.**
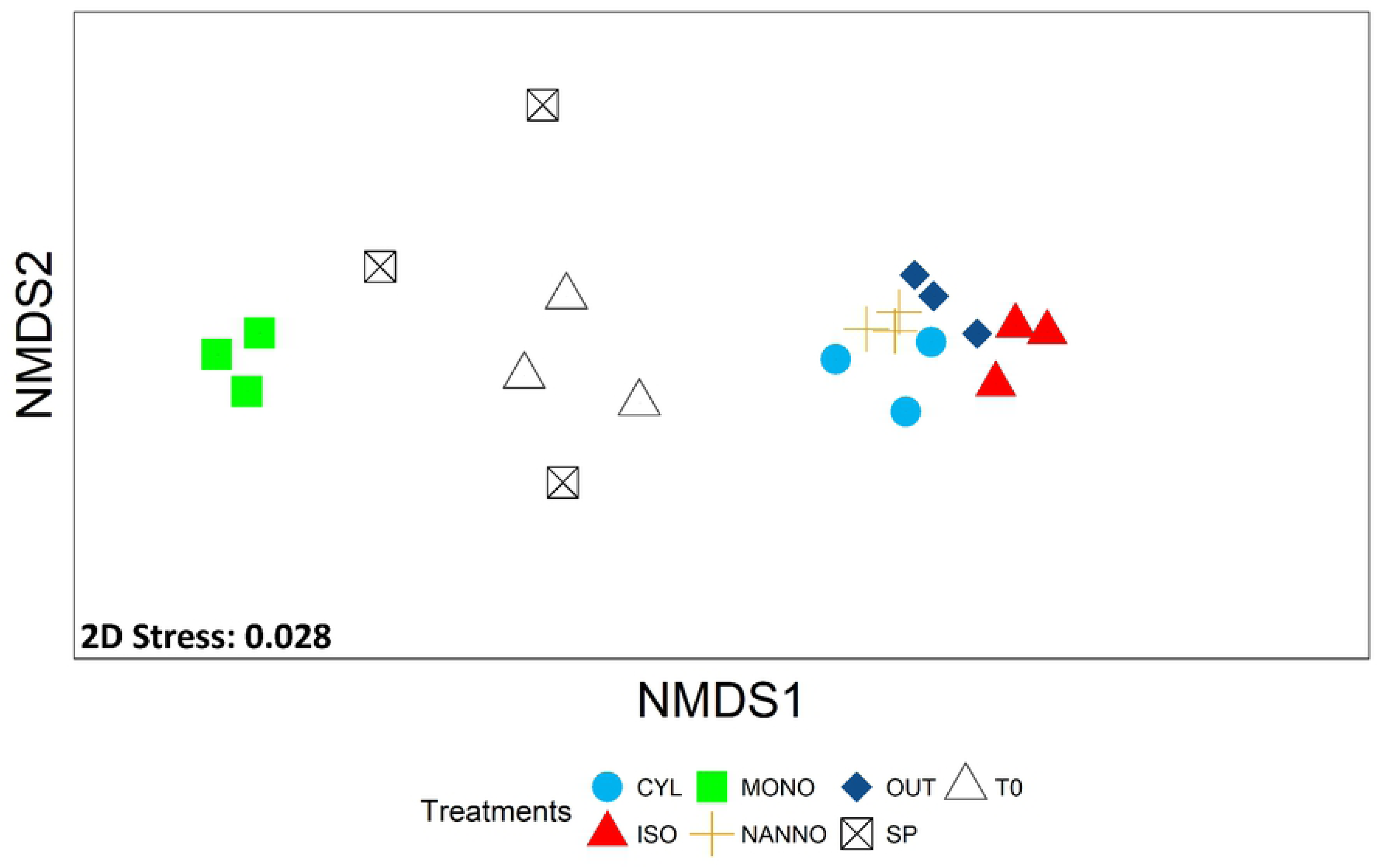
Non-metric multidimensional scaling (nMDS) plot for relative lipid class composition of the spat (%TLE) subjected to the feeding trial. T0: Spat sampled before the beginning of the trial; SP: spat fed with Shellfish Paste during the 4 weeks trial; CYL: spat fed with *C. fusiformis* 1017/2 during the 4 weeks diet trial; *ISO:* spat fed with *I. galbana* 927/1 during the 4 weeks diet trial; MONO: spat fed with *M. subterranean* 848/1 during the 4 weeks trial; NAN: spat fed with *N. oceanica* 849/10 during the 4 weeks trial; OUT: spat deployed outdoor and sampled after 4 weeks. Three replicates (n=3) for each sample group are here reported.

In absolute terms, TG resulted to be the main component of neutral lipids, and the only lipid class being affected by the diet treatments (p<0.05). The remaining neutral lipid classes were hardly affected by the diet treatments, as their absolute content remained stable between T0 and the spat diet groups (p>0.05). Indeed, the absolute content of polar lipids was also hardly affected by the different diets (p>0.05), with only LysoPC resulting significantly higher in MONO (p<0.01).

The spat lipidome exploration was completed by untargeted lipidomics. The LC-MS analysis of spat lipidome subjected to the different diets resulted in the identification of 463 features (343 of them successfully identified according to exact mass) and 620 features (267 successfully identified according to exact mass) respectively in positive (POS) and negative (NEG) ionization modes. For convenience, we will here focus on successfully identified lipid species. Plots obtained with the full dataset are available as Supplement material **(Figures S9-S12).** The LC-MS profiles and the PIT are provided in **S7 Figure** and **S1 Data**, whilst raw data are available from Mendeley data repository (DOI: 10.17632/w57zy87s68.1). POS and NEG data were separately analysed via PLS-DA, the resulting plots are shown in ***Figure 4***. Similarly to what observed on FA and lipid class analysis, the different groups showed distinct lipid profiles which resulted in neat clustering of the samples. The score plots shown in ***Figure 4*** explained the 77.6% and 58.5% between components 1 and 2 of PLS-DA models respectively for NEG and POS dataset. Cross-validation of the PLS-DA model identified in the first 4 PLS components the best model accuracy for POS and in 5 PLS needed for NEG (**Figures S6-S8**). Therefore, to extract the most meaningful for the two models we considered the average of VIP scores for the first 4 PLS components (POS) and first 5 PLS in NEG data. Lipids resulting in an average VIP scores >1 (and significantly different between treatments)are reported in ***Figure 5***.

**Figure 4:**
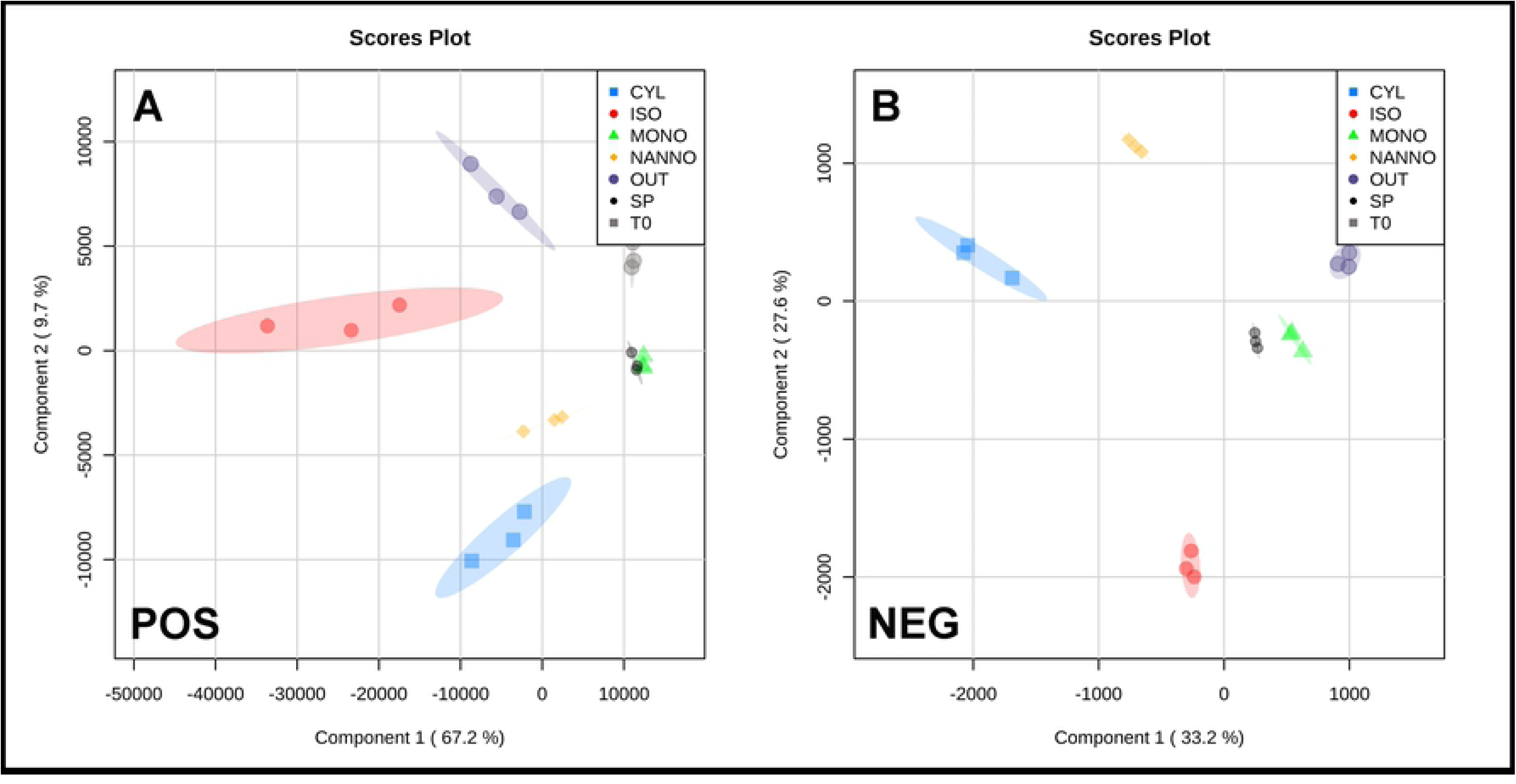
Partial least squares discriminant analysis (PLS-DA) plots of untargeted lipidomics data of the spat diet treatments groups. Positive ionization mode (A) and Negative ionization mode (B). Plotted via MetaboAnalystR.

**Figure 5.**
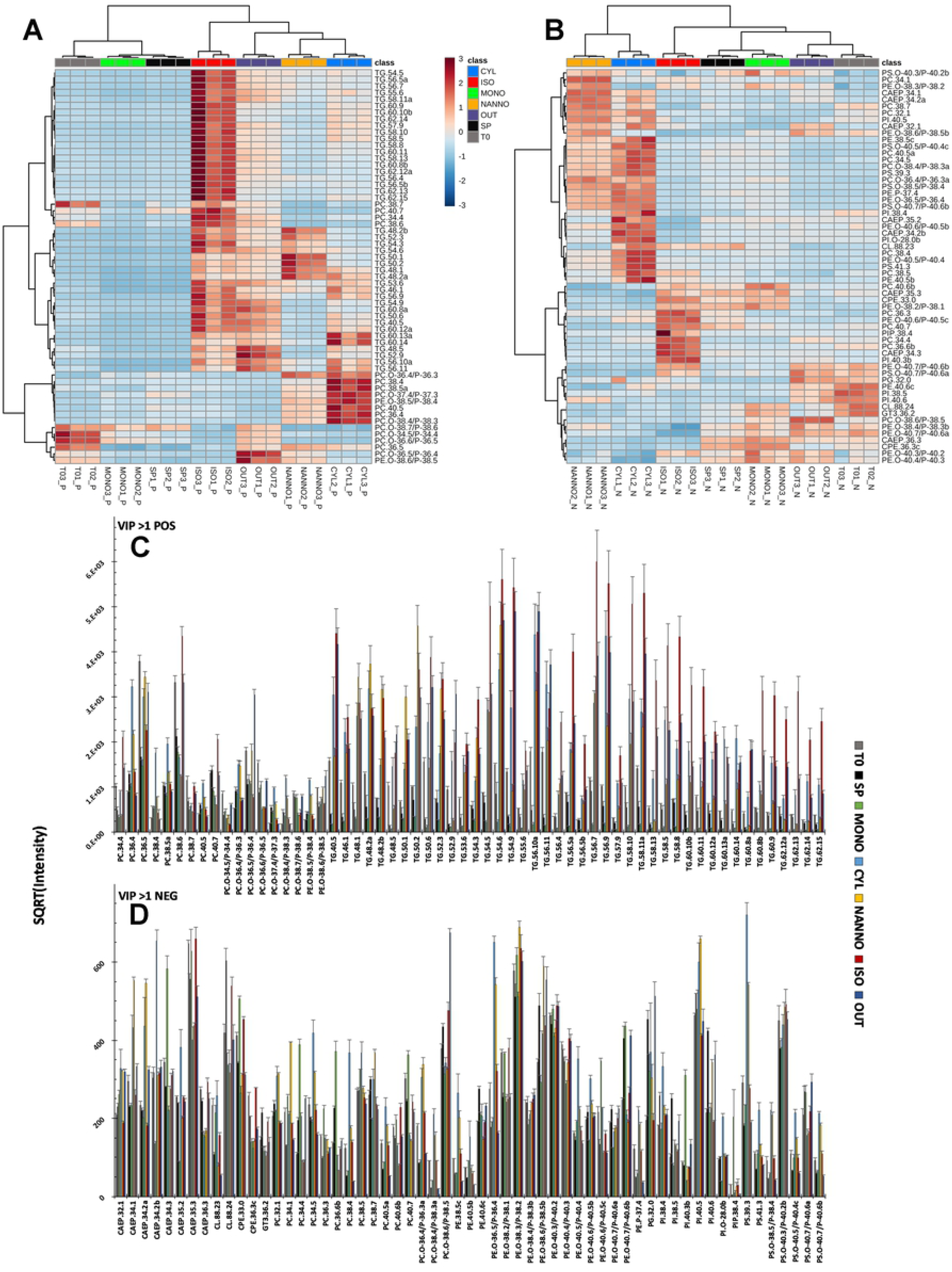
Heatmap plots and intensity charts for the VIP>1 scores evidenced by PLS-DA analysis of spat untargeted lipidomics data in Positive (A) and Negative (B) ionization mode. Euclidean distance was used as a distance measure, Ward as clustering algorithm. Lipids are reported as class, n° carbon and n° of double bonds (e.g. TG.58.10). Colour coding for lipid expression from Blue (Low) to red (High). (C) histogram plot showing the intensity of important lipids for PLSDA classification POS data. (D) histogram plot showing the intensity of important lipids for PLSDA classification NEG data. Square-rooted transformation of intensity was employed to plot the data. Data plotted via Daniel’s XL toolbox for Microsoft Excel.

In ***Figure 5A*** are reported the VIP>1 evidenced by PLS-DA analysis of POS spat data. Lipids observed in POS mode included mainly PC, PE and neutral lipids as cholesteryl esters (CE), Cer and TG. The main separation between samples in POS mode was given by the presence of TG, resulting low in T0, MONO and SP (with the remaining two overlapping in POS PLSDA plot, see ***Figure 4***). TG were abundant in the remaining groups. Unsaturated TG (n° double bonds ≥5) were principally observed in ISO treatment, while highly unsaturated TG (n° double bonds >11) resulted more abundant in CYL and ISO than in all other groups. Low unsaturated TG, as TG(48:2)a and TG(50:2), discriminated NAN from CYL, OUT and ISO diet groups. Further differences between CYL, NAN, OUT and ISO were observed in the abundance of the main PC species in each group (**Figure 5C**). PC(38:6) resulted in the most abundant PC in ISO whereas PC(35:6) was the main PC in T0, NAN and OUT. On the contrary, PC(36:4), PC(38:4) and PC(38:5)a were principally contained in CYL diet group. PC plasmalogens, PC(O-38:6/P-38:5) and PC(O-36:5/P-36:4), charaterised in OUT and in minority T0.

The heatmap plot reported in ***Figure 5B*** shows the patterns of the VIP features highlighted by PLS-DA analysis of NEG data. Polar lipid species as PE, ceramide phosphonoaminoethyl ethanolamines (CAEP), ceramide phosphonoethanolamines (CPE), PI, PS and CL are easily visualised in negative ionization mode. If compared with POS data, the patterns evidenced in NEG data are less defined (***Figure 5D***). Samples are principally clustered in two groups: one including NANNO and CYL, whilst the second one included the remaining sample groups. NANNO was characterised by high levels of PC(O-36:4), CAEP(34:1), CAEP(34:2)a and PI(40:5); PE(38:5)b, PE(P-36:4), PC(38:5)a-b, PC(38:4) and CAEP(34:2)b were highly abundant in CYL. From the other main cluster OUT was characterised by the presence of plasmalogens PC(PC(O-38:6) PC(P-38:5) and PC(O-36:5) PC(P-36:4); ISO from PC(38:6), PC(34:4), PE(P-40:5) and CAEP(34:3). MONO and SP differed for the presence of CL(88:24) in the former and CL(88:23) in the latter. T0 samples were characterised by the higher presence of PI(38:5) and PI(40:6) compared with the remaining groups. Few of the relevant compounds could not be identified via the database employed in this study, but as relevant for PLS-DA classification were kept and shown as unidentified features (see **S12 Figure** for details).

We also calculated the correlation coefficients between relevant lipids highlighted out of FA, lipid class and lipidomics with GR and WI. GR did not result in highly correlated lipids (Spearman R^2^ <0.8) so that is not discussed here (the complete correlation tables for GR and WI are provided in **S2 Data**). On the other hand, several lipids resulted correlated with WI (provided in ***Table 6***). The data suggest a positive correlation between accumulation of neutral lipid in spat and higher WI. TG (R^2^ = 0.86 p<0.05) and especially for unsaturated TG species (nn° double bonds ≥5) resulted significantly correlated with spat WI. Furthermore, also spat total content in n-3 PUFA (R^2^ ≥ 0.88 p<0.05), total PUFA (R^2^ ≥ 0.87 p<0.05), 18:1n-7, 18:2n-6 and 20:2n-6 resulted significantly correlated with WI.

**Table 6.** Spearman rank correlation coefficients of lipids highlighted from the dataset analysed in this study and spat live weight increase (WI). The reported p-values are adjusted for multiple comparisons [59].

| Spearman rank correlation coefficient |  |  |
| --- | --- | --- |
| Vs Live weight increase (WI) |  |  |
| Feature ID | R<sup>2</sup> | fdr adj P-value |
| TG(40:5) | 0.94 | 2.25E-05 |
| TG(54:9) | 0.93 | 4.60E-15 |
| TG(50:6) | 0.93 | 4.60E-15 |
| TG(58:8) | 0.92 | 4.60E-15 |
| TG(56:10) | 0.92 | 4.60E-15 |
| TG(58:11) | 0.91 | 4.60E-15 |
| TG(52:7) | 0.91 | 4.60E-15 |
| TG(56:5) | 0.91 | 4.60E-15 |
| TG(52:5) | 0.91 | 4.60E-15 |
| 20:2n-6 | 0.90 | 4.60E-15 |
| 18:2n-6 | 0.89 | 4.60E-15 |
| TG(56:7) | 0.88 | 4.60E-15 |
| Σ n-3 | 0.88 | 4.60E-15 |
| TG(54:5) | 0.88 | 4.60E-15 |
| Σ PUFA | 0.87 | 4.60E-15 |
| TG(58:10) | 0.86 | 4.60E-15 |
| TG | 0.86 | 4.60E-15 |
| TG(56:9) | 0.84 | 4.60E-15 |
| TG(58:5) | 0.84 | 4.60E-15 |
| TG(52:6) | 0.82 | 1.30E-04 |
| TG(60:10)b | 0.81 | 2.15E-04 |
| Σ Neutral lip. | 0.80 | 3.05E-04 |
| 18:1n-7 | 0.80 | 3.05E-04 |
| TG(60:14) | 0.80 | 3.54E-04 |

## 4 Discussion

For the first time to our knowledge, traditional lipid profiling techniques (as FA profiling and lipid class analysis) are here accompanied by untargeted lipidomics to approach bivalve physiology, in a comprehensive lipid analysis strategy. A large amount of information has been obtained in the past from FA analysis, however, as lipids are complex molecules with a large variety of structures and roles, many yet not fully understood in marine invertebrates [72], a comprehensive investigation is appropriate to reveal relevant information on marine invertebrate physiology [53]. The results of the three analytical approaches considered in this study offered unequivocal evidences regarding the effect of diets on mussel juveniles growth. From GR and WI analysis down to lipid molecular species, spat group could be easily classified between good performing (ISO, OUT), average performing (CYL, NANNO) and low performing (MONO, SP). The better were growth performances, the further away spat drifted from T0 samples.

### 4.1 Growth performances: who did grow, and who did not

Varying the microalgae composition of the diet has a profound effect on mussel spat growth performances [73]. OUT was the group characterised by the highest growth performances (SL 4.86 ± 0.68 mm and LW 75.08 ± 14.2 mg_LW_spat^−1^) during 4 weeks of outdoor deployment (***Figure 1***). Commonly, juvenile mussels, experience the fastest growth during summer in temperate areas [22, 74]. Typical summer phytoplankton communities in the Northeast Atlantic provide a variegate diet characterised by high abundance of diatoms and haptophytes, including high nutritional strains as *Chaetoceros sp.*, *Skeletonema sp.* and *Thalassiosira sp*., which might favour mussel growth [75, 76]. The variety of microalgae available on the water column to the spat was reflected on the FA composition of OUT, characterised by large quantity of 16:1n-7 and EPA (common markers of diatoms grazing), 18:4n-3 and DHA (markers of flagellates) and 18:2n-6 which is marker of plant and microalgae detritus [77].

Considering the results obtained from SI and WI, we could classify the tested diet groups based on observed GR as: “Fast growth” for ISO and OUT, which outperformed the remaining diet treatments in term of GR; “average growth” for CYL and NANNO, which showed significant increase in SL of spat during the feeding trial; and “Low growth” SP and MONO (no growth observed). ISO, CYL and NANNO were fed to microalgae strains rich in essential PUFA, DHA for *I. galbana* and AA/EPA for *C. fusiformis* and *N. oceanica* (**Table 2**). The observed trends agree with what was found in the past on different bivalve species, as providing either a source of DHA or EPA is known to be sufficient to meet spat nutritional requirements and obtain sustained growth [30, 32, 78]. Considering the FA composition of the diet provided, we can observe a preference for DHA rather than AA/EPA in mussel spat, as ISO resulted in significantly higher growth performances than the other two groups. Similar observations were made in *T. philippinarum* fed with *I. galbana*, showing faster growth compared with clams fed *T. suecica*, an EPA rich strain [79, 80]. ShellPaste is a commercial alternative to live algae, which although balanced in the main essential PUFA, resulted in limited growth performances of the spat. Even lower were the growth performances in of MONO; a possible reason for this could be the dietary lack of long-chain PUFA, as also observed in similar case of bivalve juveniles subjected to diets lacking of essential PUFA [37, 78, 80, 81].

### 4.2 Fatty acid analysis: de novo synthesis of essential PUFA and accumulation of specific FA in fast growing spat

After having evaluated the effect of the diets on spat growth, which largely varied between sample groups, we concentrated the efforts in studying the diet effect on spat lipid metabolism. FA composition analysis highlighted the importance of essential PUFA supply via the diet, as mussels’ spat evidenced low capabilities for *de novo* synthesis of essential n-3 PUFA; EPA and DHA have important physiological and structural roles, whilst MUFA are often catabolised or stored in reserve lipids [29, 32]. Total PUFA and total n-3 PUFA content resulted correlated with spat WI (Spearman R^2^>0.8 p<0.05, ***Table 6***). Those two parameters were depleted in slow-growing diet groups (SP, MONO). Although a similar PUFA content was provided with diets, only in nutritionally efficient diet treatments (ISO, CYL, NANNO, OUT) these two parameters were not depleted. This could be related with the tendency in bivalves to anabolise essential PUFA and catabolise non-essential FA as MUFA. Nevertheless, in conditions that do not ensure sufficient nutritional resources via the diet, also essential PUFA are catabolised to produce energy resulting in a decreasing of n-3 PUFA and total PUFA content (as observed in MONO and SP).

DHA has a principal structural function in bivalves, as suggested from several authors in the past [29, 31, 32, 80, 82–84]. The results obtained in this study show that DHA levels did not vary between the beginning of the trial and the diet groups ISO, MONO and OUT; whilst resulted lowered in CYL, NANNO and SP. This seems to be in accordance with what found by Caers, Coutteau (30), who observed that DHA content remained relatively stable in starved *or Dunaliella tertiolecta* (species lacking essential PUFA) fed *Tapes philippinarum* spat; in the same study, DHA was instead accumulated in animals fed DHA enriched diets, whilst it decreased in spat fed EPA rich diets.

Likewise, in the present study availability of EPA (*C. fusiformis*, *N. oceanica* and ShellPaste) resulted in a significant reduction of DHA in the spat with sustained growth. A reduction of DHA content was also observed in *T. philippinarum*, *Ruditapes decussatus* and *Ostrea edulis* spat subjected to EPA rich diets lacking DHA [27, 30, 42]. Da Costa (48) analysed the fatty acid assimilation in *C. gigas* larvae, observing a certain degree of elongation of EPA to 22:5n-3 in the absence of DHA. Our data suggest a different response to DHA unavailability in mussel spat, as 22:5n-3 was low in all the sample groups. On the other hand, 22:4n-6 and 22:5n-6, observed only in traces at T0 and absent on the diets provided, were accumulated in CYL and (less) in NANNO, reaching respectively the 1.55% (0.74±0.27 µgFA mg ^−1^) and 1.49% (0.71±0.24 µgFA mg ^−1^) of TLE in CYL. Similar patterns were also observed in *O. edulis* spat subjected to an EPA and AA rich diet [42]. Elongation and *de novo* synthesis of n-3 and n-6 FA are largely dependent on the supply of shorter FA precursors, with a general rule a larger synthesis of n-3 PUFA in respect of n-6 PUFA [85]. In this case, the accumulation of AA, supplied by *C. fusiformis*, might have provided a substrate for elongation to 22:4n-6 and desaturation to 22:5n-6 rather than the elongation of other n-3 PUFA to compensate for the lack in 22C PUFA (which was lacking from dietary inputs).

Non-methylene interrupted dienoic FA (NMID-FA) are endogenous FA characteristic of polar lipid in bivalves. Their exact role is not completely understood, however, they are found exclusively in the polar lipids and their content, in some cases, has been found to be inversely proportional to the essential PUFA supplied by the diet [86], whilst in other cases NMID FA were considered to be only partially influenced by dietary intakes [29, 80]. Our data agree with the first hypothesis, as NMID absolute abundance increased in ISO (20:2 NMID, diet lacking of 20C PUFA, with a significant decrease of 22:2 NMID), while the relative abundance of 20:2 and 22:2 NMID FA resulted significantly higher in SP and MONO.

18C and 20C PUFA and MUFA (18:1n-9, 18:2n-6, 20:2n-6, 18:3n-3 and 18:4n-3) were higher in ISO and OUT spat groups. In the past, these FA have been observed in neutral lipid fractions of bivalve spat [29, 31, 32, 87]. The inclusion of these FA in neutral lipids could explain for the observed high correlation of 18:1n-7, 18:2n-6 and 20:2n-6 with spat WI (Spearman R^2^ >0.8 p<0.05, ***Table 6***), as neutral lipids dominated the best performing spat groups.

### 4.3 Lipid class composition: TG content is higher in spat fed with efficient diets

The accumulation of neutral lipids (especially TG) mainly differentiated between efficient and poor diets, as evidenced from lipid class analysis, agreeing with previous authors on various bivalve species [29, 32, 80, 88]. Accumulation of TG in spat is connected with higher growth performances, as bivalve juveniles are known to accumulate lipid reserves during the summer to store energy for growth during the winter [25, 26, 88, 89]. This was also evidenced by the correlation observed by neutral lipid and TG content with WI (Spearman R^2^>0.8, p<0.05, ***Table 6***); similar correlations were observed in clams [32], mussel and scallops [90, 91] juveniles and larvae.

Main polar lipids classes (PE, PI, PS and PC) did not change between sample groups, as also reported by others [80]. Nevertheless, the two unidentified lipids Unkn_10.81_ and Unkn_12.05_ which resulted highly related to spat fed with *M. subterranean*, as these two lipids were principally observed in MONO. The polar lipids of Eustigmatales, as *Monodopsis sp.*, are known to be rich sources of glycolipids as monogalactosylglycerols (MGDG) and digalactosylglycerols (DGDG), with minor sulphoquinovosyldiacylglycerol (SQDG) [92]. The elution time-windows for these lipids, observed from previous studies applying a similar NP-HPLC separation [93, 94], could match with these unidentified lipids. Phospholipase activities in juvenile spat are lower than neutral lipases, and influenced by the diet [95]. Therefore, we could hypothesise that high content of such glycolipids, coupled with the low nutritional quality of *M. subterranean* 848/1, might have mediated the partial assimilation of them. Another polar lipid class highly present in MONO resulted LysoPC. LysoPC are products of PC metabolism, formed by the cleavage of a FA residual in position sn-1 or sn-2 by a phospholipase A [96]. Increasing in LysoPC could be connected with the lower nutritional value of *M. subterranean* 848/1 and the attempt to produce energy by polar lipid catabolism in the juvenile mussels. The catabolism of phospholipids was also observed on starved amphipods [97] and on crab larvae approaching metamorphosis [98].

### 4.4 Membrane lipids, over TG, are also affected by the dietary PUFA supply: untargeted lipidomics

Large differences in spat lipids were observed from FA and lipid class composition composition analysis, however, the most complete overview of the lipidome was obtained by LC-MS analysis in untargeted lipidomics. This approach led to the identification of 343 lipid species in POS and 267 in NEG according to MS’ (the complete lipid dataset is provided in **S1 Data**).

Similarly to what observed from lipid class analysis, TG was a major component in the discrimination between “fast growth”, “average growth” and T0 plus “slow growth” spat groups (***Figure 5***). Diets that promoted growth resulted in the accumulation of TG in spat, with several TG species highly correlated with spat WI (Spearman R^2^ >0.9 p<0.05, ***Table 6***). TG containing unsaturated FA (n° double bonds ≥5) were abundant in ISO and in OUT. These TG could be rich in 18C and 20C MUFA and PUFA which were observed in ISO and OUT spat. Few pieces of evidence are available on FA composition of specific lipid classes in bivalve spat. Caers, Coutteau (80) observed the accumulation of 18:2n-6 and 18:1n-9 principally in neutral lipid of *C. gigas* spat, whilst Soudant, Van Ryckeghem (87) reported the accumulation of 18:2n-6, 18:4n-3 and 18:3n-3 in neutral lipids of gonads of adults scallops (*Pecten maximus*) fed with T-Iso. Relevant is the case of TG (60:13) and TG (60:14), which were mainly observed in CYL. The elevated unsaturation and carbon content of these TG suggests the possible incorporation of long-chained PUFA, as AA which was copiously provided in *C. fusiformis* and accumulated in CYL. Long-chain PUFAs are commonly found esterified in polar lipids, however, when these are provided in excess of bivalves’ nutritional requirements, these can be accumulated into neutral lipids. Indeed, when Caers, Coutteau (29) supplied an excess of DHA to *C. gigas* spat, observed an increasing % of this FA in the neutral lipids fraction, although on standard conditions DHA was principally found in polar lipids. Lastly, TG(48:2)a and TG(50:2) were abundant in NANNO. Observing the NANNO FA profile, shorter chained MUFA were accumulated, so possibly these TG included 16:1n-7 and 18:1n-9 and SAFA as FA residuals.

Working at single lipid species level allowed to capture changes in the composition of polar lipids of the spat, which were not observed from lipid class analysis due to the different separation employed in the two techniques. PC and PE are the main polar lipid components in bivalves [62, 64, 65, 67, 87, 99]. Bivalves have well-conserved PC structures, with a SAFA (16:0 and minor 18:0) in sn-1 position and an unsaturated PUFA (EPA or DHA) on sn-2 [99]. Recently these structures have been also confirmed via LC-MS/MS on *M. edulis*, observing that the most common PC species in bivalves resulted in PC(36:5), PC(38:6) and PC(38:5) and for the plasmalogens PC(O-36:5), PC(O-38:5), PC(O-38:6), PC(P-38:5) [64]. These are mostly in agreement to what observed for T0 spat, which was characterised by PC(36:5), PC(38:6) and their relative plasmalogens PC(O-36:5/P-36:4) and PC(O-38:6/P-38:5). The plasmalogens were principally observed in the OUT samples, possibly as a result of higher water temperatures, as an increase of plasmalogens on mussels between winter/spring and summer conditions was found by Facchini, Losito (62). When spat were fed *I. galbana*, PC(38:6) was highly abundant, as this might have incorporated a DHA molecule in the sn-2 position. Lower was the content of PC(36:5), compared with T0 in this group, due to the lack of EPA obtained via the diet and observed from ISO FA profile. On the contrary, large content of PC(36:5) was observed in NANNO, as *N. oceanica* was the richest source of EPA. Whilst PC(38:4), PC(38:5)a and PC(36:4) were abundant in CYL and in a lower extent in NANNO. Both *C. fusiformis* and *N. oceanica* provided AA, which might have been employed for synthesis of PC(36:4), whilst PC(38:4) could include 22:4n-6 as PUFA and PC(38:5)a could have been constituted by 22:5n-6 (both FA accumulated in CYL).

The NEG mode was dominated by polar lipids including PE, PI, CAEP and CL (**Figure 5B-D**). PE are the second major class of bivalves phospholipids [80, 99, 100], characterised by a small ethanolamine polar head, and are observed in specific domains of cell membranes as ion channels and cell-cell connections [50]. In bivalves, a large percentage of PE (around 40%) are plasmalogens [64, 65]. The analysis of NEG data suggests that the PE with the highest intensity belonged all to the plasmalogen species (***Figure 5D***). The exact role of plasmalogens in bivalves is not fully understood, some hypothesis suggests their role as membrane permeability (as commonly rich in long-chain PUFA) and adaptation to environment changes [99]. The effect of the diet treatments on PE composition was less pronounced than what observed on PC, as a consequence of the large content of plasmalogens observed in this lipid class, which could not be resolved without the aid of MS/MS [101]. Nevertheles, PE abundance in the spat diet groups could be somehow connected with the different dietary inputs. Lipids with 1-2 double bonds characterised NANNO, which was the group that accumulated the largest amount of FA 16:0, 16:1n-7 and 18:1n-9 (**Table 5**). Whilst PE and PS with 4 and 5 double bonds were largely shared between CYL and NANNO (**Figure 5B**). PE(O-40:6/P-40:5)c was largely found in ISO and PE(38:4/P-38:3)b, PE(40:6)c, PE(O-40:7/40:6)a-b were mainly observed in T0 and OUT. SP and MONO were characterised by PE(O-38:2/P-38:1), PE(O-40:3/P-40:2) and PE(O-40:4/P-40:3).

Few PI were also relevant in group classification, with PI(40:5) abundant in CYL and NANNO, PI(40:3)b in ISO, and PI(38:5) and PI(40:6) in T0 and OUT. PI is a substrate for phospholipase C, yielding DAG (which are directed for phospholipid synthesis [102]) and inositol-1,4,5-trisphosphate (IP_3_), an important secondary messenger in Ca^2+^ channels regulation. IP_3_ is suspected to influence gonad maturation and fertilization in bivalves [104, 105]. The FA composition of PI can have a relevant role for physiological processes as reproduction and growth in bivalves. PI are commonly rich in AA and other long-chained PUFA, resulting in an important substrate for eicosanoids and prostaglandins production [99, 106].

Several CAEP species were also responsible for group clustering of NEG lipidomics dataset. CAEP is an important class of polar lipid in bivalves, often the third most abundant after PC and PE [107]. CAEP and the closely related CPE belong to ceramide lipids and represent for invertebrates the analogues of vertebrates’ sphingomyelins. Their existence and roles were mostly unknown until the last decades when have been observed in several invertebrate species ranging from jellyfish to bivalves and insects [108, 109]. The chemical stability of CAEP, given by the sphingosine backbone and the C-P bond, making them between the most refractory components of the lipidome. The higher expression of CAEP(35:3) CAEP(36:3) observed in MONO and SP could be related to their resistance to degradation and action of lipases [63].

CL is a further group of phospholipids, in which a third glycerol molecule is acetylated in position sn-1’ and sn-2’ creating a pseudo symmetrical molecule [110]. CL are predominantly located inner mitochondrial membrane where exert a relevant role in the oxidative phosphorylation process [111]. CL(88:24) and CL(88:23) were listed between the main lipids explaining for PLS-DA classification. in SP and MONO, compared with the remaining diets. Long-chain PUFA CL are common in bivalves, and the presence of CL(88:24) has been observed in the past in *P. maximus*, *C. gigas* and *M. edulis* [112], explaining for the relatively high abundance of such lipid in T0. In bivalves, CL composition is highly conserved in each between species, while tend to diverge between different molluscs [113, 114]. However, stress conditions, resulting in oxidative stress and ROS production, also influence CL composition, due to the function of this particular lipid in cellular respiration [115]. Other authors evaluated the dietary effect on mitochondrial CL, observing an increase of PUFA containing CL in mice subjected to SFA and MUFA rich diets, and on the contrary, a relative decrement of certain PUFA containing CL in presence of PUFA rich diets [116]. This trend is interestingly similar to what observed in MONO and SP, which were subjected to nutritionally poor (SP) and PUFA low (MONO) diets. Nevertheless, no final conclusion on this observation could be made, as CL(88:23) was also highly observed in CYL and ISO (subjected respectively to EPA/AA and DHA rich diets).

## 5 Conclusions

Nursery bivalve production is the most economically demanding of bivalve hatchery practices, and at the same time newly settled juveniles are extremely delicate, suffering of high mortality right at the end of the hatchery production cycle, when their product value is at its maximum. Therefore, expanding existing knowledge of the nutritional physiology of this key life stage will have significant effects for mussel aquaculture production and the ecology of this important species. Mussel spat represented an interesting model organism, as coupled fast metabolic rates (observed in the large differences observed between diet treatments in the relatively short time-frame of 4 weeks) and a simple holding set up (more robust and resistant of mussel larvae) to responses that are directly comparable to adult mussels.

Among the tested diets, *I galbana* offered the best results in term of GR and WI between laboratory-reared spat, suggesting the importance of DHA source for efficient growth of young mussels. The combination of the three lipid analysis techniques provided valuable information on spat lipid metabolism. Classical lipid profiling techniques as FA and lipid class analysis permitted easily to discriminate between low-performing and high-performing spat groups. Providing at least one source of essential C20 or C22 PUFA was necessary to the spat, resulting in minimal or absence of growth when no C20/C22 PUFA were supplied (MONO). Our data also suggest that the supply of only one main PUFA (EPA or DHA) might stimulate the spat to counterbalance the lack of the other essential FA (e.g. CYL subjected to surplus of AA, elongated to 22:4n-6 and desaturating that to 22:5n-6 and in ISO, with increasing in content for FA 20:3 NMID under EPA shortage). FA analysis also evidenced how the best groups growth-wise, OUT, ISO, CYL and NANNO, accumulated MUFA, 18C and 20C PUFA during the feeding trial. These groups were also characterised by large TG content as evidenced by lipid class analysis, which differentiated them from T0, SP and MONO.

All the jigsaw pieces were connected by lipidomics, which offered a clearer image of spat lipidome, (together) with the joint use of the other two traditional lipid analysis approaches. The advantage of lipidomics compared with traditional lipid profiling approaches dwells on the study of lipid molecular species. Through lipidomics, we observed a TG specific accumulation patterns between diets, linked with the accumulation of specific MUFA and PUFA in the spat. Furthermore, lipidomics evidenced deep changes in the relative abundance of main membrane lipid species in relation to the dietary essential PUFA supply; trends which were missed by lipid class profiling. A reason for this, aside the higher sensitivity of the LC-MS apparatus, links with changes related to shifts in lipid molecular species abundance for each polar lipid class, rather than actual changes in lipid class abundance, and this is also the reason why, to our knowledge, it is the first time that such pattern is reported on bivalve juveniles. Visualising this trend through traditional lipid analysis techniques would require the FA profiling of purified lipid classes samples, often not feasible due to limited sample availability and instrument sensitivity. Lipidomics is powerful to streamline and expand lipid analysis possibilities on marine organisms, however, it comes with large and convoluted data, requiring the necessary ability to cope with that.

In the future, lipidomics could be a tool to answer several important biological questions regarding lipid metabolism of marine invertebrates, while at the same time expanding our knowledge on lipid classes which present logistical difficulties when tackled with traditional lipid profiling techniques. So far, the coverage of lipid ID available for marine invertebrates is rather small (74% for POS and 43% for NEG lipids could be confidently identified using existing online lipid mass spectra libraries), and acquisition of further fragmentation spectra is necessary to expand such online resources. The advancement of LC-MS platforms is an important driver for this process, but only the continuous application of these techniques on marine organisms can expand our possibilities to clearly define their metabolism and responses to the environment.

## Acknowledgements

The authors declare no conflict of interest with the topic presented in this paper. We would like to thank Inverlussa Marine Services (www.inverlussa.com) for their support and spat supply and the Culture and Collection of Algae and Protozoans (CCAP) for the supply of the algae strains. The authors thanks also the Nutrition Analytical Services of the Institute of Aquaculture of the University of Stirling, Mr. James Dick for support and expertise during FAME analysis.

## Captions to supplementary material

### Figures

**S1_Fig: Turbidimetry data recorded before and after every feeding of the spat.**

**S2_Fig: Liquid chromatography mass spectrometry (LC-MS) profiles of spat total lipid extracts (TLE).** ESI POS mode profiles left traces, NEG mode profiles right traces. Data are aquired at precursor ion MS (MS’) via high resolution LC-MS platform (Exactive, ThermoScientific). Plotted via Excalibur 4.1 (ThermoScientific).

**S3_Fig: Principal Component Analysis of POS lipidomics spat dataset Plotted via MetaboAnalystR.** Cylindrotheca: CYS, Isochrysis: ISO, Monodopsis: MONO, NannoChloropsis: NANNO, Outdoor: OUT, QC: Quality control samples, Shell Paste: SP, T0: T0 samples.

**S4_Fig: Principal Component Analysis of NEG lipidomics spat dataset Plotted via MetaboAnalystR.** CYS *Cylindrotheca fusiformis* fed spat, ISO *Isocrysis galbana* fed spat, MONO *Monodopsis subterranean* fed spat, NANNO *Nannochloropsis oceanica* fed spat, OUT Outdoor deployed spat, QC: Quality control samples, SP *ShellPaste* fed spat, T0: T0 samples.

**S5_Fig: PLS-DA analysis of spat lipidomics dataset POS Model fitting: 2000-fold Permutation test.** Plotted and calculated via MetaboAnalystR.

**S6_Fig: PLS-DA analysis of spat lipidomics dataset POS: Model fitting: 10-fold leave one out –Cross-validation analysis LOOCV.** Q_2_ used as parameter of model fitting. Plotted and calculated via MetaboAnalystR..

**S7_Fig: PLS-DA analysis of spat lipidomics dataset NEG: Model fitting: 2000-fold Permutation test.** Plotted and calculated via MetaboAnalystR.

**S8_Fig: PLS-DA analysis of spat lipidomics dataset NEG: Model fitting: 10-fold leave one out – Cross-validation analysis LOOCV.** Q_2_ used as parameter of model fitting. Plotted and calculated via MetaboAnalystR.

**S9_Fig: Partial least squares discriminant analysis (PLS-DA) plots of untargeted lipidomics data acquired in positive ionization mode.** Full data is used in this model including unknown features. *Plotted via MetaboAnalystR*.

**S10_Fig: Partial least squares discriminant analysis (PLS-DA) plots of untargeted lipidomics data acquired in Negative ionization mode.** Full data is used in this model including unknown features. *Plotted via MetaboAnalystR*.

**S11_Fig: Heatmap plot for the top 30 VIP evidenced by PLS-DA analysis of spat untargeted lipidomics data in Positive ionization mode.** Full data is here used, including unknown features. Euclidean distance was distance measure, Ward as clustering algorithm. Lipids are reported as class, n° carbon and n° of double bonds (e.g. TG5810). Colour coding for lipid expression from Blue (Low) to red (High).

**S12_Fig: Heatmap for the top 30 VIP scores evidenced by PLS-DA analysis of spat untargeted lipidomics data in Negative (NEG) ionization mode.** Euclidean distance was used as distance measure, Ward was used as a clustering algorithm. Lipids are reported as class, n° carbon and n° of double bonds (e.g. TG.58.10). Colour coding for lipid expression from Blue (Low) to red (High). In absence of an exact mass ID features are reported as ret. time_mass/charge (e.g. 8.86_1019.5664m/z) or ret. time_neutral mass (e.g. 11.13_1330.7461n).

### Tables

**S1_Table: Main adducts and exact masses (for positive and negative ionization) of the lipid standard mixture used for exact mass (MS’) identification of lipidomics data.**

### Data

**S1_Data: Peak intensity tables obtained by the processing of lipidomics data via Progenesis QI software.** During data processing chromatograms were aligned to a QC samples, peak picking was done following automatic configuration of the software and inserting a threshold background noise of 1xE5 (POS) and 1xE4 (NEG). Samples are reported as normalised abundance. The reported PIT was furtherly processed via MetaboanalysR as reported in the main text of the paper. Compounds identifiers are found on Column A. Samples are found after column G. Accepted identification codes are given according Lipid Maps and HMDB codes. Lipid ID was provided from exact MS’ with Δppm <5 and relative adduct (as reported in S1_Table).

**S2_Data: Spearman correlation matrix between important lipids highlighted from fatty acids (FA), lipid class and lipidomics (POS/NEG) dataset and growth rate (GR) and weight increase (WI) of spat.** Relative abundances were employed for lipidomics data, whilst absolute values were used for FA and lipid class. Correlation values were calculated via R statistical software (v 3.5).

